# Choice-driven remapping of action- and stimulus-anchored value in human single neurons

**DOI:** 10.64898/2026.05.28.728546

**Authors:** Aniek Fransen, Cooper D. Grossman, Chrystal M. Reed, Jeffrey M. Chung, Adam N. Mamelak, Ueli Rutishauser, John P. O’Doherty

## Abstract

Whether stimulus- and action-based choices rely on a shared neural architecture remains intensely debated. While lesion and fMRI studies show conflicting regional dissociation and overlap, the underlying single neuron substrates remain unknown. We recorded single neuron activity from the human ventromedial prefrontal cortex (vmPFC), anterior cingulate cortex (ACC), pre-supplementary motor area (preSMA), and amygdala during distinct action- and stimulus-based reward tasks. Prior to decision-making, neurons across all regions broadly encode value for both choice domains. However, choice commitment triggers domain segregation across the prefrontal cortex. While ACC and vmPFC neurons flexibly track both chosen actions and stimuli, they exhibit a double dissociation for chosen valuation: ACC neurons selectively maintain action-based values, whereas vmPFC neurons selectively maintain stimulus-based values. Conversely, preSMA neurons invariant to task domain track chosen action identity and value. These findings reveal how prefrontal networks transition from domain-general evaluation to domain-segregated choice execution.

## Introduction

Value-based choices rely on the computation of expected values for potential candidates, whether those candidates are visual stimuli or motor actions (Rangel et al., 2008). A fundamental debate in the study of value-based decision-making is whether these value computations are segregated by the choice-domain (i.e. stimulus versus action-based choices) or instead rely on a shared neural substrate (see ‘common currency’ literature e.g., Levy and Glimcher, 2012). Historically, specific neural circuits have been theoretically partitioned based on these choice-domains. For instance, evidence from lesion studies in both monkeys and humans have demonstrated a macro-level functional dissociation, revealing specific action or stimulus selection deficits following damage to the anterior cingulate cortex (ACC) and orbitofrontal cortex (OFC), respectively (see Rudebeck et al., 2008; Camille et al., 2011). While these findings suggest a broad regional divide in necessary neural infrastructure, it remains unclear if such distinct separation appears at the cellular level. Additionally, these studies leave open the question of how these action- and stimulus-based value signals are dynamically computed over the stages of the decision process.

To characterize the encoding of stimulus and action values, previous studies have employed various recording modalities, though critical gaps remain. Single-neuron studies in non-human primates have been instrumental in mapping valuation circuitry, frequently identifying regions such as the striatum, premotor cortex, and anterior cingulate cortex as key nodes for action-based decision-making (e.g., Samejima et al., 2005; Scangos and Stuphorn, 2010; Hayden and Platt, 2010; Seo and Lee, 2007; Kennerley et al., 2006). However, a pervasive methodological limitation in much of this literature is the inherent conflation of motor actions with spatial targets. In some experimental paradigms, animals are required to indicate their choices by executing a movement (such as a saccade or a reach) to a specific spatial coordinate (e.g., the left or right side of a computer screen). As such, the motor actions performed only varied with respect to the spatial location they were directed towards. Consequently, neural signals that have been interpreted as encoding ‘action value’ might simultaneously reflect the value of the spatial target, the spatial attention directed toward it, or the directional movement vector itself (see for discussion; Padoa-Schioppa, 2011). This confound makes it exceedingly difficult to isolate ‘pure’ motor action value. Here we therefore aimed to define an action space that spanned unique physical effectors (e.g., selecting a left-hand versus a right-foot movement) and was as dissociated from the visual stimuli/attention as possible, in an effort to more directly pair value to distinct motor actions.

In parallel to investigations of action value, a robust line of single-neuron research has sought to characterize the encoding of stimulus value, primarily focusing on the orbitofrontal cortex (OFC) and ventromedial prefrontal cortex (vmPFC). Studies in non-human primates have demonstrated that neurons in these regions encode the economic value of offered and chosen visual stimuli, often doing so independently of the sensorimotor contingencies required to obtain them (Padoa-Schioppa and Assad, 2006; Strait et al., 2014). This has led to the prominent hypothesis that the OFC/vmPFC computes a purely abstract, domain-general “common currency” for economic choice, evaluating choice options independent to any motor planning (Padoa-Schioppa and Assad, 2006; Levy and Glimcher, 2012). However, testing whether these stimulus value signals are truly distinct from, or integrated with, action value computations requires directly recording from both domains simultaneously within the same neural populations.

In the human brain, functional magnetic resonance imaging (fMRI) has been deployed to investigate whether these valuation domains are represented by distinct networks. Such studies have yielded evidence for regional overlap, implicating the ventromedial prefrontal cortex (vmPFC) in encoding value-related signals supporting choices over stimuli and actions (Gläscher et al., 2009; Wunderlich et al., 2010). Additional studies have set out to map the process of stimulus to action mapping in stimulus-based decisions and found a network involving vmPFC, dorsomedial PFC and motor cortex to support such stimulus-to-action decisions (Hare et al., 2011). However, while fMRI has successfully mapped relevant valuation circuitry, its reliance on blood-oxygen-level-dependent (BOLD) signals inherently limits both spatial- and temporal resolution. A single functional voxel exhibiting overlapping representations of stimulus and action value could reflect a genuinely integrated, mixed-selective neural population, but it could equally arise from entirely segregated (but anatomically interleaved) populations of domain-specific neurons. Similarly, the poor temporal resolution of the hemodynamic response function makes differentiating between distinct facets of the decision process difficult. fMRI activity related to value signals within a trial could reflect neuronal firing pre-choice, post-choice or a mixture (or compounding) of both. Temporally segregating the neural response to each of these decision stages is necessary to gain a deeper mechanistic understanding of these stimulus- and/or action-based value computations. Direct electrophysiological recordings in humans are uniquely positioned to bridge these gaps and resolve the question of true single-neuron separability.

Previous research utilizing human single-unit recordings has successfully identified neurons that represent the value of available choice candidates, regardless of whether they are chosen (here also referred to as “decision value”), in pre-supplementary motor area (preSMA) during value-based choice (Aquino et al., 2023). Yet, because the behavioral task employed in this study was not designed to decouple the visual stimuli used from the physical motor effectors required to select them, it remains unclear whether these human value signals are stimulus- or action-based in their format, or part of a domain-general computation.

To address this open question, we set out to differentiate the encoding of stimulus value and action value at the single-neuron level in the human brain. We designed an experiment to tackle the two limitations discussed above. We included a stable set of stimuli and actions, allowing us to track the expected value attached to each of these choice options throughout the experiment. Randomization of stimulus-action pairing additionally orthogonalized stimulus- from action values. Lastly, we employed clearly distinct motor actions and tied reward probabilities to these motor responses in the action portion of the experiment. This set of hand- and foot-operated buttons helped isolate action value as it is tied to different motor effectors. We recorded single-unit activity from 902 neurons across the vmPFC, ACC, preSMA, and amygdala to investigate the neural representation of stimulus and action value. We leverage the temporal resolution of these recordings to trace the computation of value as the decision process evolves from the pre-decision representation of choice alternatives to choice commitment and selection.

## Results

Our primary objective was to characterize and compare the single-neuron computations underlying value-based decision-making when choices are framed over motor actions versus visual stimuli. To address this, we recorded 230 vmPFC, 299 preSMA, 157 dACC and 216 amygdala single neurons in 20 sessions across 16 patients implanted with hybrid macro- and micro-electrodes for epilepsy monitoring. Participants performed an experiment specifically designed to differentiate the encoding of stimulus- and action-based decisions. As part of this experiment, participants preformed two structurally parallel reinforcement learning tasks in counterbalanced order (see Methods for details). In the “Action Task,” reward probabilities were tied to specific motor responses. Given our questions’ emphasis on distinct action value encoding, we chose to enrich the action-space beyond the typical 2 hand-button press used in previous choice experiments. As such, our motor actions included a left-hand button press, a right-hand button press, a left-foot pedal press, and a right-foot pedal press. In contrast, in the “Stimulus Task,” reward probabilities were associated with visual stimuli. The Stimulus Task took advantage of the same four motor actions for choice selection. Importantly, we ensured the value of visual stimuli was independent of the physical action required to select them through trial-wise pseudo-random stimulus-action pairing (see Figure 1A, B and Methods). Participants completed over five blocks of 36 choice trials on each task on average (see Methods).

**Figure 1:**
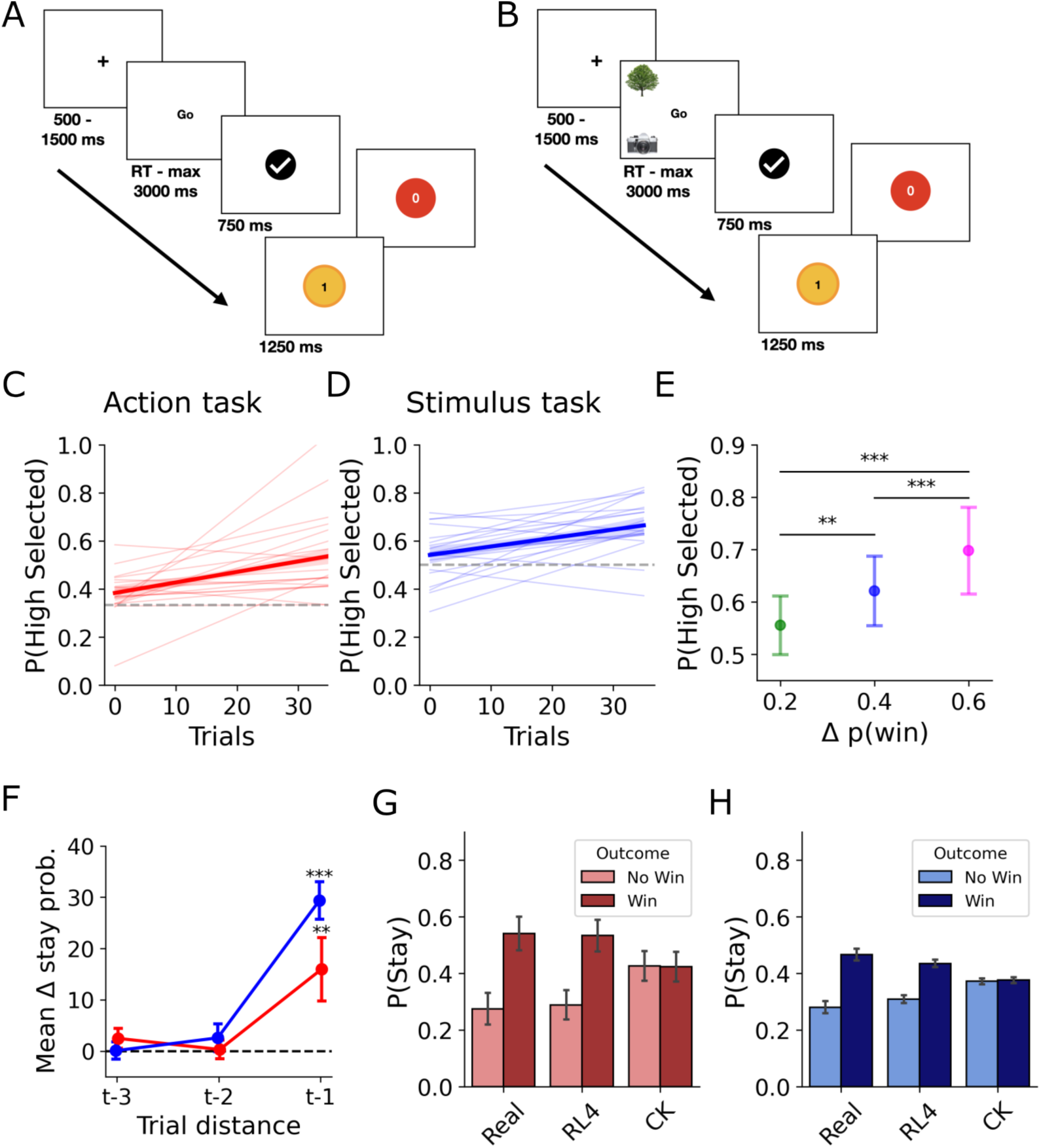
Task and behavioral results. See below for continued caption. **A)** Trial design of action task. Fixation cross is followed by the “Go” signal to indicate one of three instructed actions can be chosen and executed. Actions include; left hand press, left foot press, right hand press and right foot press. The check mark confirms action selection independent of the specific action chosen. The trial ends with either a reward (yellow coin) or no reward (red coin) in accordance with the reward probability attached to the chosen action. Participants complete 36 such trials over the course of a block. See Methods for more details on task design. **B)** The stimulus trial design mirrors the action task with the exception that stimuli are displayed in the “Go” window. There are four total stimuli; tree, house, dog and camera. Participants make a choice between the 2/3 pseudo-randomly drawn stimuli displayed on the screen. The stimulus location determines the action required to make the selection. Stimuli displayed on the top of the screen can be selected using the hand buttons (i.e. left hand to select the tree in example) while stimuli on the bottom of the screen can be selected using their left or right foot (i.e. left foot to select camera in example). In this task the probability of winning (i.e. reward) is dependent on the stimulus selected (and independent of the action executed). **C)** Probability of selecting the candidate with the highest expected value. Solid line indicates average performance across participants and blocks in the action task (with the shaded band representing the 95% confidence interval). Individual session’s performance is shown in the thinner background lines. **D)** Average performance similarly increases over the course of blocks in the stimulus task. Please note chance here is P=1/2 since participants select one out of two available candidates in each trial. **E)** Probability of selecting the correct candidate (i.e. highest expected value) in the stimulus task increases with the difference in win probability between the available candidates. In other words, participants perform better on trials that are easier. Whiskers indicate standard error around the mean (indicated with the central dot). These effects are significant (Mann-Whitney U test: ** : *p <* 0.01 and *\*\*\** : *p <* 0.001) **F)** Participants are more likely to repeat an choice that was rewarded on the previous trial (t-1) across both tasks. We see this effect quickly decays further back into the past where at two trials (t-2) and three trials back (t-3) this effect is no longer significant. Whiskers indicate standard error around the mean. Estimates from a logistic regression model (win vs no win, see Methods) transformed to log odds probabilities. **G** The probability of staying after a win on the previous trial (darker colored bars) or a no-win on the previous trial (lighter colored bars) in the action task. Bars indicate the mean across sessions with the whiskers representing the standard error. Leftmost bars display real participant stay tendencies. The middle bars are simulated using the RL model (see Methods), while rightmost bars indicate choice kernel model simulations (see Methods). **H** Model simulated stay behavior in the stimulus task. Plot follows the same conventions as G.

### Behavioral Results

We started our investigation of participants’ behavior across these two tasks by determining if participants were able to learn to select more rewarding choice candidates (i.e., actions in the Action Task and stimuli in the Stimulus Task, note that *options* is a more typical word used for decision candidates but we refrain from doing so as to avoid confusion with the notion of an *option* from reinforcement-learning (Sutton et al., 1999)). Participants could learn to select candidates with a higher probability of reward through experiencing the win or no-win outcomes resulting from the selection. As such, we would expect to see an increase in the selection of the most rewarding choice candidate over the course of a block. Indeed participants demonstrated significant learning over the course of a 36-trial block in both tasks. In the Action Task, the probability of selecting the candidate with the highest expected value increased significantly over time (Wilcoxon test over the last 10 trials: *p <* 0.001; Figure 1C). At the start of a block, when the reward probabilities associated with each action are unknown, participants’ performance was at chance level. However, this performance improved over the course of the block indicating participants learned to associate the available actions with their reward probabilities (see Methods). Similarly, in the Stimulus Task, participants selected the more rewarding stimulus more than expected by chance at the end of a block (Wilcoxon test over the last 10 trials: *p <* 0.001; Figure 1D).

Furthermore, choice accuracy in the Stimulus Task was sensitive to the relative value of the available stimuli. The probability of selecting the most rewarding choice candidate available increased monotonically with the difference in win probability (Δ*p*(*win*)) between the two presented stimuli (Figure 1E). This indicates that participants were more effective at identifying the optimal choice when the expected value difference was larger (i.e., when the choice was easier).

#### Temporal Dynamics of Reinforcement

To investigate how recent reward history shaped choice behavior, we performed a session-level logistic regression analysis to determine the probability participants repeat rewarded choices (Figure 1F and see Methods). We hypothesized that if participants were utilizing reinforcement to guide their behavior, they would be more likely to repeat a choice that resulted in a win. Our results confirmed this predictions across both tasks. We observe participants are approximately 15 to 30% more likely to repeat the choice for which they received a reward on the immediately preceding trial (for action and stimulus tasks respectively; *p* = 0.009 for action and *p <* 0.001 for stimulus task). Notably, this effect is highly similar across both tasks, suggesting that the integration of rewards follows a similar temporal scale regardless of whether the reward is associated with a motor action or a visual stimulus.

#### Computational Modeling of Choice Behavior

Finally, we sought to formalize participants’ choice behavior by fitting a computational model. We fit a reinforcement learning (RL4) model and a choice kernel (CK Only) model to the participants’ choice data (see Methods for model details). The choice kernel model captures participants tendency to repeat previously chosen actions (see Figure 1G, H). However, this model is unable to replicate the observed difference in selection repetition after a win versus no-win (see Supplemental Figure S2 for quantitative model comparison results). We therefore utilized a four-parameter reinforcement learning model that tracks both expected values (*Q*-values) and choice history (choice kernel), allowing us to capture both value-based updates and choice perseveration (see Figure 1G,H and Supplemental Table S2 and Figure S2).

To connect our behavioral findings with the hypothesized single-neuron computations, we utilized the selected reinforcement learning model to extract latent, trial-by-trial decision variables for each session. While our behavioral analysis demonstrated that this model successfully captures the dynamics of choice, its primary utility for our neural analysis lies in generating precise, session-specific estimates of expected values. This process allows us to isolate these internal computational signals as they evolve over the course of a block through experience with the outcomes (wins or no-wins). We can then use these latent variables as direct trial-by-trial regressors to elucidate the associated neural computations. As a result, we can map exactly where and when individual neurons across our regions of interest dynamically track these specific decision signals during action- and stimulus-based choices.

### Neural representations of Action and Stimulus Decision Value

Having established that participants utilize a reinforcement-based strategy in both tasks, we turn to investigating the underlying neural computations. Our primary goal was to determine whether single neurons in our areas of interest (vmPFC, preSMA, ACC and amygdala) represent value across choice domains (i.e., coding for the value of both stimuli and actions) or if these representations are domain-specific. There are various points in a trial where we might hypothesize different types of value signals are computed. We start by analyzing the representation of the value of choice options that are available for selection. These value signals will serve as an input to the comparison/decision process. Here we will refer to these values that proceed choice as ‘decision value’ (note that these types of value signals are sometimes referred to as ‘offer value’ (e.g., Shi et al., 2022; Fine et al., 2023)).

Based on previous literature (Camille et al., 2011; Kennerley and Walton, 2011), we might hypothesize domain-specific value computations, with regionally clustered neurons in the prefrontal cortex specializing in either action- or stimulus-based decision value representation. To test this hypothesis, we identified neurons whose firing rates significantly correlated with the decision value of actions or stimuli available for selection, as derived from our RL model, after stimulus onset. In the action task, we surprisingly found a significant population of neurons across vmPFC (*n* = 46, 20%, *p <* 0.001), preSMA (*n* = 46, 15.4%, *p <* 0.001), ACC (*n* = 35, 22.3%, *p <* 0.001) and amygdala (*n* = 30, 13.9%, *p <* 0.001) that tracked the value of available actions (Figure 2B; proportions evaluated via binomial tests on permuted neuron significance, see Methods). A temporally resolved encoding analysis (see Methods) additionally highlighted that neurons across these areas encode the expected value (*Q*-value) of available actions early, even before the ‘Go’-screen prompting choice initiation.

**Figure 2:**
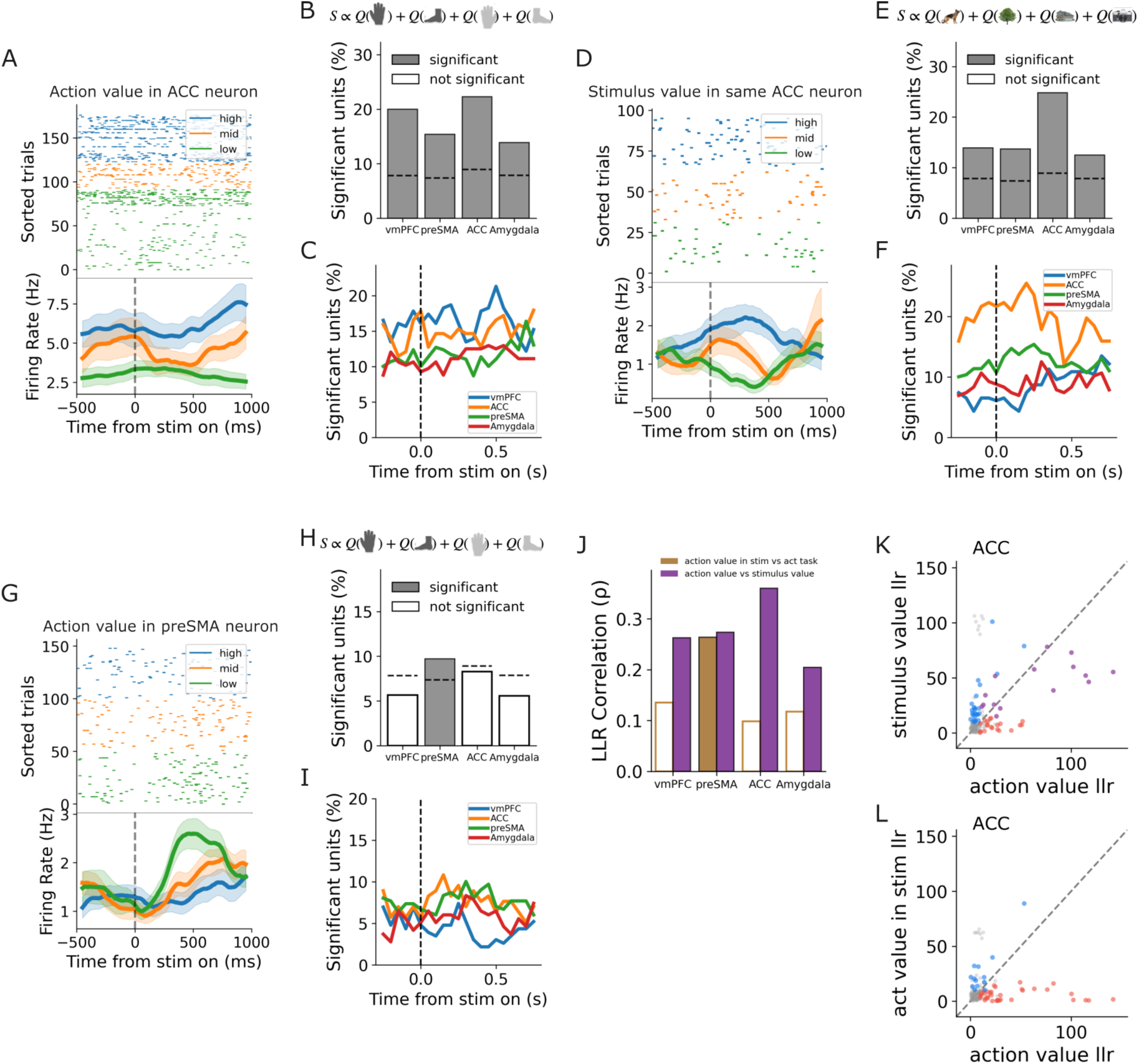
Decision value coding. See below for continued caption. **A)** Action value representation in an example Anterior Cingulate Cortex (ACC) neuron. Top: Raster plot of individual trials aligned to stimulus onset (dashed vertical line at 0 ms). Trials are sorted and color-coded by action value tertile (blue: high, orange: mid, green: low). Bottom: Peristimulus time histogram (PSTH) showing the average firing rate for each tertile. Shaded regions represent the standard error. **B)** Proportion of significant neurons encoding action value during the action task across four recorded brain regions: ventromedial prefrontal cortex (vmPFC), pre-supplementary motor area (preSMA), ACC, and Amygdala. The generalized linear model used for estimation is displayed above, highlighting the four available action options. Dark gray bars represent the percentage of significant units, while white bars represent the remaining non-significant population. Dashed lines indicate the chance level. **C)** Time-resolved proportion of significant units encoding action value relative to stimulus onset, calculated using a sliding window approach (see Methods). Solid lines denote the proportion for each brain region at *p <* 0.05 uncorrected. **D)** Stimulus value representation in the exact same ACC neuron shown in panel A, demonstrating mixed selectivity. The raster plot and PSTH follow the same conventions as in panel A, but trials are sorted by stimulus value tertiles during the stimulus task. **E)** Proportion of significant neurons encoding stimulus value across the four brain regions. The corresponding stimulus value model is shown above, highlighting the distinct visual stimuli. **F)** Time-resolved proportion of significant units encoding stimulus value relative to stimulus onset, following the same conventions as panel C. **G)** Action value representation in an example preSMA neuron, following the same conventions as panel A. **H)** Proportion of significant neurons encoding action value during the stimulus task, following the conventions of panel B. **I)** Time-resolved proportion of significant units encoding action value during the stimulus task relative to stimulus onset. **J)** Correlation in model statistics across tasks and value domains. Bar plot displays the Spearman rank correlation (*ρ*) of likelihood ratio statistics for each brain region. Dark orange bars compare action value coding in the stimulus task versus the action task. Dark purple bars compare action value coding versus stimulus value coding across the respective tasks. Filled bars indicate the correlation is significant at *p <* 0.01. **K)** Scatter plot comparing the model statistic for stimulus value in the stimulus task (y-axis) against action value in the action task (x-axis) for individual units in the ACC. Units are colored based on significance across both tasks. We then label neurons red when they are significant only in the action task, blue if they solely contribute significantly in the stimulus task. Units labeled in purple if they contribute significantly across both tasks/variables. Dashed line added to the 45-degree line for visual aid in comparing units to ‘equal contribution’. **L)** Scatter plot comparing the model statistic for action value in the stimulus task (y-axis) against action value in the action task (x-axis) in the ACC. Conventions follow panel K. No units are found to be significant across both tasks.

Similarly, in the stimulus task, a significant population of neurons across the regions of interest tracked the value of the available stimuli (vmPFC: *n* = 32, 14%, *p <* 0.001; preSMA: *n* = 41, 13.7%, *p <* 0.001; ACC: *n* = 39, 24.8%, *p <* 0.001; amygdala: *n* = 27, 12.5%, *p <* 0.001)(see Figure 2E). In elucidating the temporal dynamics of these signals, we observed a clear stimulus-onset-induced ramping of encoding (Figure 2F). In the vmPFC, specifically, we observed a steady increase in the number of neurons that code for stimulus-based decision value as time progresses post stimulus onset.

Lastly, we evaluated the representation of available action values during the stimulus task. Although reward probabilities in this task were explicitly tied to the visual stimuli, the brain must ultimately translate these expected values into specific motor plans to execute a decision. To test whether the brain represents pre-decision action values in a stimulus-based task, we mapped the expected value of each presented stimulus to the specific motor effector required to select it on that given trial (e.g., if a stimulus yields a Q-value of 0.8 and its current paired action is the left-hand button press, the available left-hand action value is set to 0.8). This operationalization, which is standard for computing action values in spatial stimulus-based bandit tasks (e.g., Aquino et al. (2023)), enables us to test whether the brain automatically evaluates potential motor actions even when the decision is primarily stimulus-driven. This analysis revealed significant action decision value encoding solely in the preSMA (preSMA: *n* = 29, 9.7%, *p <* 0.001; vmPFC: *n* = 13, 5.7%, *p* = 0.37; ACC: *n* = 13, 8.3%, *p* = 0.052; amygdala: *n* = 12, 5.6%, *p* = 0.40 - see Figure 2H).

Together, these results reveal that, contrary to our hypothesis, there is substantial overlap in the regions that encode for stimulus decision value in the stimulus task and action decision value in the action task. However, this does not yet determine whether the same neurons underpin these computations across domains. We tested this domain generality at the single-neuron level by computing the stability of model statistics of single neurons across tasks (Figure 2J; see Methods). Here we observe a significant correlation between action value and stimulus value coding strength across tasks in all ROIs (vmPFC: *ρ* = 0.26, *p <* 0.001, preSMA: *ρ* = 0.27, *p <* 0.001, ACC: *ρ* = 0.36, *p <* 0.001, amygdala: *ρ* = 0.20, *p* = 0.003, see Figure 2J in purple). These correlations indicates that neurons strongly encoding the value of actions in the action task were significantly more likely to also encode the value of stimuli in the stimulus task. Conversely, neurons that contributed to action value coding in the stimulus task were not more likely to code for action value in the action task. This pattern appears in all areas except preSMA (*ρ* = 0.26, *p <* 0.001, see 2J in orange) and is particularly striking in the ACC (ACC: *ρ* = 0.10, *p* = 0.22, vmPFC: *ρ* = 0.14, *p* = 0.04, amygdala: *ρ* = 0.12, *p* = 0.08). We thus conclude that the neural representation of decision values is supported by a more overlapping set of neurons across choice-domains than the representation of action values per se. In other words, neurons are more likely to encode the decision value in whichever domain is relevant to the current choice than they are to maintain a domain-specific encoding regardless of task relevance (i.e., always code for action value).

In the ACC we can clearly observe this dichotomy when plotting the individual neurons’ model statistics (colored by significance; Figure 2K, L). Among ACC neurons, we can observe four distinct clusters: a subpopulation coding for both stimulus and action value (purple), alongside strictly action-selective (red), strictly stimulus-selective (blue), and non-value-coding (gray) populations. This is in stark contrast to the cross-task action value comparison (Figure 2L). While we observe neurons that code for action in each task there is a complete lack of overlap in these populations.

When further investigating the coding scheme employed in the subset of neurons that code for both stimulus values and action values (e.g., those in purple in Figure 2K) we find a remarkably consistent flip in coding directionality in ACC neurons (see Figure S3). Nearly all neurons in the ACC found to encode stimulus value do so using a positive coding scheme (where the firing rate increases with value). In contrast, these same neurons in the ACC largely negatively code for action value in the action task. While the number of neurons in this analysis is small this result might point to the ACC using anti-correlated subspaces to encode decision values across these different choice domains. Together, these results demonstrate a common neural substrate for domain-relevant decision values (while relying on different subspaces within the ACC particularly).

### Representations of Chosen Value

Having detailed the neural encoding of decision values, we next investigated whether these regions also reflect the value of the final decision (i.e., the chosen value). More specifically, we sought to determine if the expected value of the selected choice candidate (chosen *Q*-value) is represented across our ROIs and whether this encoding differs depending on the task domain. Again, previous research might lead us to hypothesize a regional specialization in either stimulus- or action-value. Alternatively, there could be a continued domain-flexibility as the choice progresses from decision value to chosen value.

To differentiate between these possible hypotheses, we evaluated chosen value encoding during the Action Task. In the period leading up to the response, we found that chosen action values were selectively represented in regions heavily implicated in motor planning and action valuation. Specifically, a significant proportion of neurons in the preSMA (*n* = 31, 10.4%, *p <* 0.001) and the ACC (*n* = 17, 10.8%, *p* = 0.002) tracked the value of the action ultimately executed by the participant (see Figure 3A, B). In contrast, we did not find significant encoding of chosen action value at our predefined threshold (*p <* 0.01) in the vmPFC (*n* = 17, 7.4%, *p* = 0.071) or the amygdala (*n* = 18, 8.3%, *p* = 0.024).

**Figure 3:**
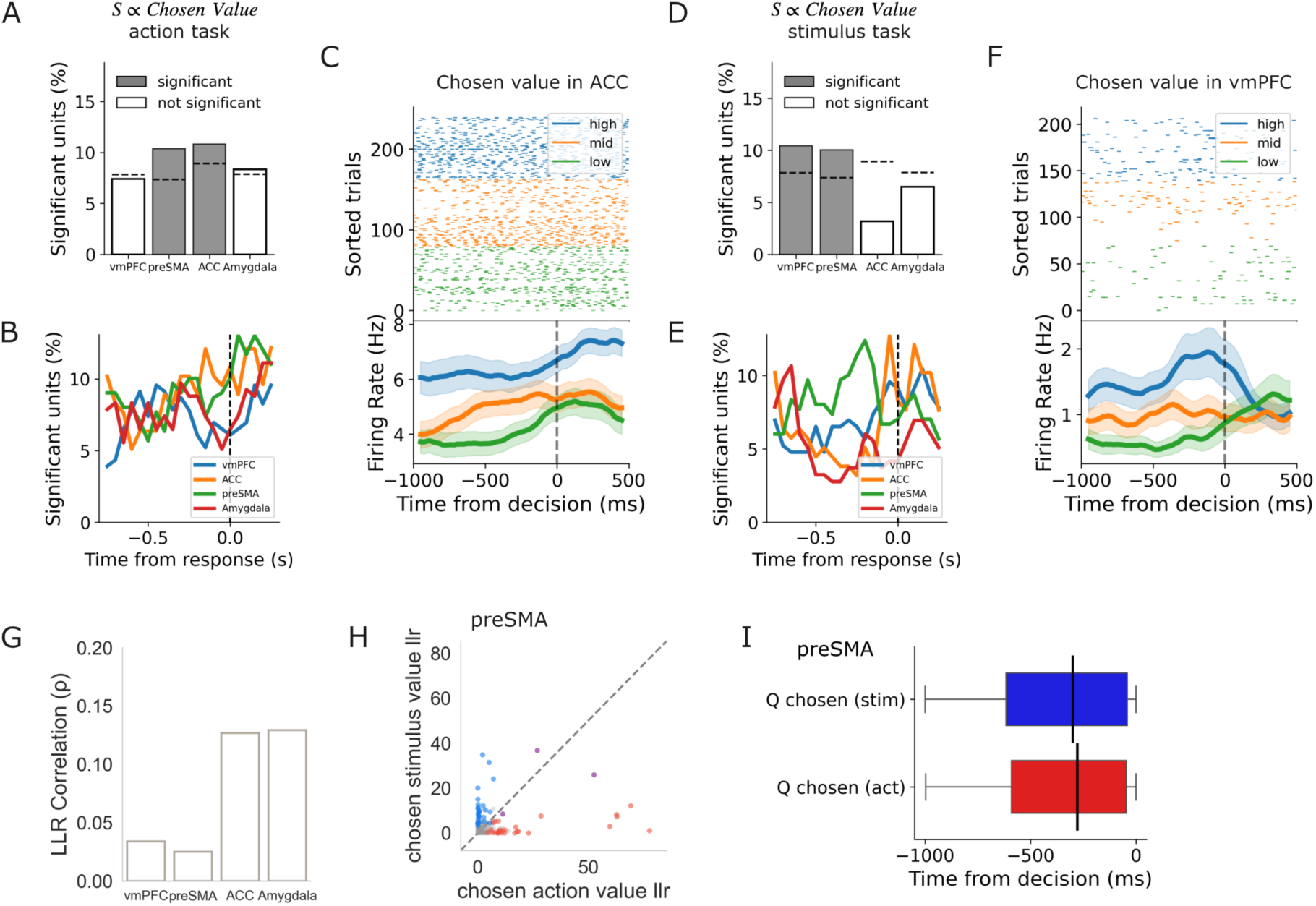
Chosen value encoding. See below for continued caption. **A)** Proportion of significant neurons encoding chosen value during the action task across four recorded brain regions: ventromedial prefrontal cortex (vmPFC), pre-supplementary motor area (preSMA), anterior cingulate cortex (ACC), and Amygdala. Dark gray bars indicate regions where the percentage of significant units exceeds chance (at *p <* 0.01), while white bars represent non-significant populations. Dashed horizontal lines indicate the specific chance level for each region (see Methods). **B)** Time-resolved proportion of significant units encoding chosen value relative to response execution (dashed vertical line at 0 ms) in the action task. Solid lines denote the proportion for each brain region computed using a sliding window approach (see Methods). **C)** Chosen value representation in an example ACC neuron during the action task. Top: Raster plot of individual trials aligned to response onset. Trials are sorted and color-coded by chosen value tertile (blue: high, orange: mid, green: low). Bottom: Peristimulus time histogram showing the average firing rate for each tertile. Shaded regions represent the standard error. **D)** Proportion of significant neurons encoding chosen value during the stimulus task. Conventions are identical to panel A. **E)** Time-resolved proportion of significant units encoding chosen value relative to the response during the stimulus task. Conventions follow panel B. **F)** Chosen value representation in an example preSMA neuron during the stimulus task. Conventions follow panel C. **G)** Spearman correlation for chosen value across tasks. The bar plot displays the correlation (*ρ*) of chosen value model statistics between the stimulus task and the action task for each brain region. None of the correlations are significant. **H)** Layout as in Figure 2K. Scatter plot comparing the model statistic for chosen value in the stimulus task (y-axis) against chosen value in the action task (x-axis) for individual units in the preSMA. Units are colored based on significance across both tasks (red for significance in action task; blue for the stimulus task; purple for both; gray for neither). Dashed line added to the 45-degree line for visual aid in comparing units to ‘equal contribution’. **I)** Latency analysis of the onset of firing rate change in significant units. Significance determined based on comparison with the expected Poisson baseline firing rate (see Methods). Analysis aligned to response (indicated by time 0).

The same analysis run on the Stimulus Task revealed a dissociation in chosen value coding. Rather than utilizing a static, domain-general network across both decision domains, we observed a partial shift in the regions recruited to represent chosen stimulus value. In the stimulus domain, the vmPFC reliably encoded the value of the chosen option (*n* = 24, 10.4%, *p <* 0.001), alongside the preSMA (*n* = 30, 10.0%, *p <* 0.001). Conversely, the ACC (*n* = 5, 3.2%, *p* = 0.897), which had prominently tracked chosen value in the action task, failed to exhibit significant coding for chosen stimulus value, as did the amygdala (*n* = 14, 6.5%, *p* = 0.195) (see Figure 3D, E). To formally test this double dissociation between the vmPFC and ACC, we utilized a logistic regression to evaluate the likelihood of a neuron significantly encoding chosen value as a function of region and task (see Methods). This analysis revealed a highly significant region × task interaction (*β* = 1.68, *z* = 2.72, *p* = 0.006), confirming a double dissociation.

Together, these results highlight a more specialized, domain-specific set of neurons track the value of the chosen option in stimulus- versus action-framed choices. Unlike in the encoding of pre-choice decision values, at the region-level we already observed a domain-specific specialization. A significant proportion of neurons in the ACC track the chosen value exclusively when decisions are made over motor actions. Conversely, the vmPFC demonstrates the opposite specialization, being selectively recruited to track chosen value only during stimulus-based decisions.

Bridging this domain-specific dichotomy is the preSMA, which encodes chosen value regardless of the choice domain. To further characterize the encoding of chosen value at the level of individual neurons, we tested whether a neuron more strongly encoding chosen value in the stimulus-domain would equally encode chosen value when the choice was action-driven (correlation of model statistics - see Methods). This analysis revealed that the encoding of chosen value in stimulus- and action-domains relies on separated sub-populations (vmPFC: *ρ* = 0.03, *p* = 0.61, preSMA: *ρ* = 0.02, *p* = 0.67, ACC: *ρ* = 0.13, *p* = 0.11, amygdala: *ρ* = 0.13, *p* = 0.06, see Figure 3G). This separation of value coding appears even in the preSMA, where we did observe both stimulus and action chosen value at the region level (see Figure 3A, D, G, H). From this analysis we conclude that different subsets of neurons within the preSMA facilitate the representation of domain-specific chosen value codes. Maintaining such information would be relevant for mapping to motor plans and credit-assignment (see Discussion). Lastly, despite being driven by separate sub-populations in the preSMA, an analysis of the latency of chosen value coding revealed no latency difference between the encoding of chosen values in the stimulus and action task (see Figure 3I). As such, while these chosen values are encoded in regionally segregated neurons, the similar temporal evolution of this computation might hint at its shared function across choice-domains.

### Representations of Chosen Identity

Having established that distinct regional populations specialize in tracking the value of the chosen candidate depending on task domain, we next investigated whether these regions also represent the identity of the chosen candidate. Note that an isolated value signal is insufficient for goal-directed behavior; the brain must explicitly encode the identity of the chosen candidate to successfully orchestrate its selection and to properly assign credit during subsequent learning updates. We would therefore hypothesize that the regions found to represent chosen value would also represent the chosen identity.

We first evaluated the chosen action identity (the specific button press) encoding across our regions of interest during the action task. Remarkably, we observed robust representation of the selected action across prefrontal regions. Specifically, a highly significant proportion of neurons in the preSMA (*n* = 56, 18.7%, *p <* 0.001), the ACC (*n* = 41, 26.1%, *p <* 0.001), but also the vmPFC (*n* = 24, 10.4%, *p <* 0.001) tracked the identity of the chosen action (see Figure 4A). A temporally resolved analysis further revealed that the proportion of engaged neurons in both the ACC and preSMA exhibits a ramping trajectory that progressively builds as the moment of choice execution approaches (see Figure 4B). Such ramping would be consistent with the hypothesis that these signals might contribute to the motor planning leading up to selection. The amygdala, however, did not exhibit significant encoding of chosen action identity in the action task (*n* = 15, 6.9%, *p* = 0.126).

**Figure 4:**
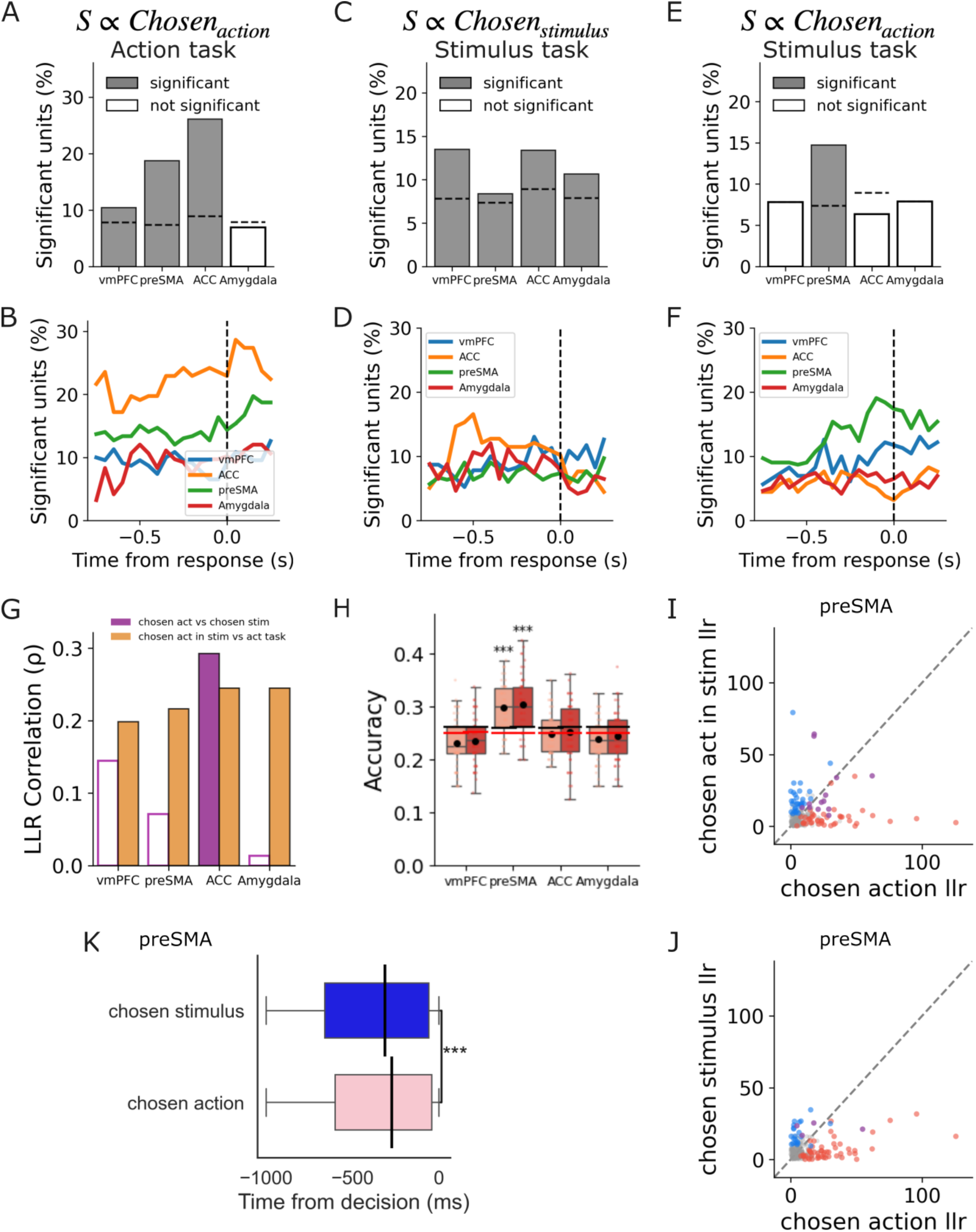
Chosen identity coding. See caption below. **A)** Proportion of significant neurons encoding the chosen action (*Chosen_action_*) during the action task across four regions of interest: ventromedial prefrontal cortex (vmPFC), pre-supplementary motor area (preSMA), anterior cingulate cortex (ACC), and Amygdala. Dark gray bars indicate regions where the percentage of significant units exceeds chance (at *p <* 0.01); white bars represent non-significant populations. Dashed horizontal lines indicate the region-specific chance levels. **B)** Time-resolved proportion of significant units encoding the chosen action relative to response execution (dashed vertical line at 0 ms) in the action task. Solid lines denote the proportion for each brain region computed using a sliding window approach (see Methods). **C)** Proportion of significant neurons encoding the chosen stimulus (*Chosen_stimulus_*) during the stimulus task. Conventions follow panel A. **D)** Time-resolved proportion of significant units encoding the chosen stimulus relative to the response during the stimulus task. Conventions follow panel B. **E)** Proportion of significant neurons encoding the chosen action during the stimulus task. Conventions follow panel A. **F)** Time-resolved proportion of significant units encoding the chosen action relative to the response during the stimulus task. Conventions follow panel B. **G)** Correlation between model statistics for chosen identity across tasks. The bar plot displays the Spearman rank correlation (*ρ*) comparing unit model statistics for chosen action versus chosen stimulus within the stimulus task (purple), and chosen action in the stimulus task versus the action task (orange) across brain regions. Filled bars indicated a significant *p <* 0.01 correlation while outlined bars are non-significant correlations. **H)** Generalization performance of chosen identity decoding across brain areas: vmPFC, preSMA, ACC, and Amygdala. Box plots show cross-condition decoding accuracy for different train-test combinations. Lighter red boxes show decoding performance for models trained on action identity in the chosen action task and tested on the chosen action identity in the stimulus task. Darker red boxes show the reverse: models trained on the chosen action identity in the stimulus task and test on the action task. Asterisks denote decoding performance significantly above the chance level (*\*\*\** : *p <* 0.001). The chance threshold (*p <* 0.05) is indicated with a black line while the mean of the permuted chance distribution (see Methods). **I)** Layout as in Figure 2K. Likelihood ratio statistic for individual units in the preSMA for either chosen action in the stimulus task (on y-axis, with significance in blue) or chosen action in the action task (on the x-axis, with significance in red). There are a few neurons that code for chosen action significantly across both tasks (labeled in purple). **J)** Likelihood ratio statistics of chosen stimulus (in stimulus task, in blue along the y-axis) and chosen action (as in Figure 4I). **K)** Latency analysis for preSMA units significantly coding for chosen stimulus identity or chosen action identity in the stimulus task. Latency distributions compared using two-sided Wilcoxon rank-sum test (*\*\*\** : *p <* 0.001). Analysis run aligned to response (indicated by time 0).

We next examined chosen action identity during the stimulus task. In this task, the motor output is merely a vehicle to obtain a desired visual stimulus. Crucially, under these conditions, the comprehensive representation of the to-be-executed action collapsed in most regions. At our predefined threshold (*p <* 0.01), the preSMA was the only region that maintained a significant representation of the motor response (*n* = 44, 14.7%, *p <* 0.001). Strikingly, the ACC (*n* = 10, 6.4%, *p* = 0.261) and the vmPFC (*n* = 18, 7.8%, *p* = 0.042) ceased to reliably track the identity of the action, as did the amygdala (*n* = 17, 7.9%, *p* = 0.045). This result replicates findings from previous research (e.g., Aquino et al. (2023)) that found the preSMA to encode chosen action identity in a stimulus-based choice task. Additionally, the clear ramping pattern observed in preSMA (see Figure 4F), further reinforces the interpretation that this area might translate value information to motor commands to relay to neighboring motor cortex. When comparing this encoding of chosen action identity across tasks, we find a subset of preSMA neurons significantly codes for chosen action identity across both tasks (see Figure 4G, I, preSMA: *rho* = 0.22, *p <* 0.001, vmPFC: *ρ* = 0.20, *p* = 0.003, ACC: *ρ* = 0.24, *p* = 0.002, amygdala: *ρ* = 0.24, *p <* 0.001). Additionally, a generalization analysis in this area confirms this motor action representation is indeed task invariant in the preSMA (see Figure 1H and Methods).

In contrast to this specialized preSMA encoding of chosen action in the stimulus task, the encoding of chosen stimulus identity appeared more widespread. During the response period of the stimulus task, we found widespread, significant encoding of the chosen stimulus identity across all of our recorded regions of interest. The vmPFC (*n* = 31, 13.5%, *p <* 0.001), ACC (*n* = 21, 13.4%, *p <* 0.001), amygdala (*n* = 23, 10.6%, *p <* 0.001), and preSMA (*n* = 25, 8.4%, *p* = 0.009) all possessed significantly large neural populations dedicated to signaling which visual stimulus the participant had committed to (see Figure 4C). While we do also observe significant encoding of chosen stimulus in the preSMA, this code depends on a distinct sub-populations of neurons than that for chosen action in the action task (see Figure 4J & G). An analysis of the onset of these chosen identity codes in the stimulus task shows the chosen stimulus encoding proceeds the chosen action encoding in the preSMA (see Figure 4K). This finding supports the idea that the preSMA might translate stimulus information into action selection.

Lastly, while there is a significant correlation between single neuron encoding strength across choice domains in the ACC (*ρ* = 0.29, *p <* 0.001), the encoding of chosen stimulus identity and action identity appear to rely on independent sub-groups of neurons within vmPFC, preSMA and amygdala (vmPFC: *ρ* = 0.14, *p* = 0.03, preSMA: *ρ* = 0.07, *p* = 0.22, amygdala: *ρ* = 0.01, *p* = 0.84, see Figure 4G in purple). We can thus conclude that while there is regional overlap in the encoding of chosen identity across task-domains, this encoding is distinctly more segregated than we observed for decision value encoding (see Figure 2J).

The ubiquity of these chosen identity signals might hint at their function in maintaining the code necessary to ensure the new value assigned to the chosen candidate after outcome gets appropriately assigned. However, our limited set of recording locations prevents characterizing these processes in, for example, the basal ganglia. The temporal resolution of these recordings does however offer us a potential for a post-hoc test to falsify the role of these chosen identity signals in such an outcome-driven update tagging. If these representations are indeed used in value-assignment, we might expect to see persistent representation continuing into the outcome window. Alternatively, if the encoding of chosen identity is temporally limited to the response period this would exclude the possibility of their use for tagging an outcome-driven value update.

We thus extended our analysis of chosen identity to the outcome window (see Supplemental Figure S4). This analysis revealed robustly maintained encoding of chosen actions in the Action Task and chosen stimuli in the Stimulus Task across prefrontal ROIs (see Supplemental Figure S4A, C). When plotting the time course of these signals we find strong persistent encoding of the chosen action into the outcome window particularly for the ACC and preSMA in the action task (see Supplemental Figure S4B). For the stimulus task this temporally resolved analysis shows maintained encoding (particularly) in vmPFC and what appears as re-instantiation in the ACC and preSMA (see Supplemental Figure S4D). This finding stands in contrast to the encoding of chosen action in the stimulus task, for which we observe a decline in encoding prevalence post choice execution (see Supplemental Figure S4E, F). This temporally restricted encoding to the response-window confirms that chosen action coding in the stimulus task is being used exclusively for choice execution. In contrast, the observed pattern in domain-relevant chosen identity coding is consistent with what we would hypothesize if these signals fulfilled a role in updated-value assignment (see Discussion).

## Discussion

An extensive debate has revolved around whether stimulus- and action-based choices rely on a shared neural architecture, with lesion studies and human fMRI work appearing to find opposing evidence (e.g., Rudebeck et al., 2008; Gläscher et al., 2009). By recording from single neurons across a distributed prefrontal network during both action- and stimulus-based decisions, our results unify these findings by suggesting that the brain relies on a dynamic architecture within which the representations of action and stimulus value reorganize as the decision unfolds. Prior to choice commitment, valuation recruits broad, partially overlapping populations of neurons across the prefrontal cortex across both choice-domains. However, upon commitment to a choice, these populations undergo a remarkable functional bifurcation, separating into highly specialized, modality-specific codes.

In the pre-decision valuation phase neurons across the prefrontal cortex (including both the vmPFC and ACC) code the decision value for both actions and stimuli. These representations of action- and stimulus-based value use a shared neural substrate, where the a neuron coding for one value-domain is more likely to also code for the other. We want to briefly note however, that this does not prove the existence of a true ‘common currency’ across these choice domains (Padoa-Schioppa, 2011; Levy and Glimcher, 2012). While we do find a small subset of neurons that similarly encode action- and stimulus value, there are also preliminary signs of a domain-specific value coding scheme within neurons in the ACC. Future research will have to probe the population-level geometry of these value representations to determine if these value computations indeed rely on (semi-) orthogonal subspaces (as suggested (although strictly at the population-level) in Johnston et al., 2024). These findings resonate with human fMRI literature reporting overlapping BOLD activations for both stimulus and action values in prefrontal cortex regions (Gläscher et al., 2009). Crucially, our temporally resolved single-unit data reveals that any overlapping encoding is a strictly pre-decision phenomenon.

As the decision process transitions to the commitment phase we observe a more segregated, modality-specific architecture. This suggests the prefrontal value encoding network is actively reshaping its representations to separately encode the value of chosen actions and stimuli. Such a regional specialization across choice-domains could facilitate the correct encoding of the “value-relevance” of either the action and stimulus. Inappropriately assigning the outcome to, for example, the action chosen in a stimulus-based decision could be detrimental for value learning. Indeed, segregating the representation of chosen action values from chosen stimulus values would allow the brain to directly link selected candidate to the relevant domain (as might be necessary for credit assignment see below). Reflecting this need for compartmentalization, we observe that domain-overlap ends during the post-choice valuation phase, revealing a stark double dissociation between vmPFC and ACC. The ACC specializes by selectively maintaining action-based chosen value representations. This finding extends classical non-human primate literature (e.g., Hayden and Platt, 2010) by demonstrating that this specialization applies even to more clearly separated motor actions. Conversely, the vmPFC selectively maintains stimulus-based chosen value representations, supporting its role as a post-choice evaluator of stimuli (Wunderlich et al., 2010; Hare et al., 2011). This network-level dichotomy provides a compelling neuronal account for macro-level lesion studies that reported specific action or stimulus selection deficits following damage to the ACC and OFC, respectively (Rudebeck et al., 2008; Camille et al., 2011).

Additionally, in the commitment phase, we find the prefrontal cortex broadly represents the identity of the chosen option across both action- and stimulus-based choices. For example, both the vmPFC and ACC maintain this cross-task encoding for choice identity (dynamically tracking the chosen action or chosen stimulus) but they do so utilizing largely independent sub-populations within these regions. Beyond immediate choice execution, this post-decision representation of chosen identity and value within the relevant choice domain is necessary for effective credit assignment in value-based learning. When an outcome (i.e., win or no-win) is received after choice, the brain must determine whether to attribute that outcome to the physical action taken or the visual stimulus chosen. We hypothesize that by selectively isolating the chosen action value within the ACC and the chosen stimulus value within the vmPFC, the brain establishes dedicated structural channels as it prepares the input to an outcome-based value update. Such an architecture would allow prediction error signals to selectively update the value of the relevant choice dimension.

We are, however, limited in our observation of this update process due to our clinically defined recording locations. Direct signals of such value updates have been found in lateral PFC, basal-ganglia and the midbrain (e.g., Valentin and O’Doherty, 2009; Asaad and Eskandar, 2011; Lak et al., 2014; Man et al., 2024). Despite this limitation, we do find a widespread and persistent chosen identity coding that would be consistent with these signals fulfilling a secondary function beyond correct choice execution. The chosen identity’s function in selecting the relevant option would only require it to be presented until that selection is complete. Indeed, while we observe this post-response ramp down for chosen action identity in the stimulus task, the task-relevant chosen identity persists into the outcome window. Such a persistent encoding would be expected if the chosen identity additionally facilitates a tagging mechanism linking an updated value to the correct item. Future research should aim to further elucidate the updating of both action- and stimulus-based value signals. Ideally these studies will be able to directly record from areas of basal-ganglia in conjunction with a wide set of prefrontal regions (including lateral PFC). A combined recording across these regions will be able to expand our analysis of value signals across choice-domains to additionally include reward-based updates.

Outside of the various valuation signals found in the vmPFC and ACC, our results position the preSMA as a highly dynamic encoder of both stimulus and action information. Rather than acting as a simple final motor output region, the preSMA exhibits highly flexible encoding of pre-decision value, chosen value and chosen identity regardless of the choice domain. Furthermore, we observed a distinct temporal cascade of single-neuron activity within the preSMA during stimulus-based decisions. Specifically, the encoding of the chosen stimulus identity significantly precedes the encoding of the chosen action. This sequential encoding mirrors observations in the supplementary eye field (Chen and Stuphorn, 2015) and strongly aligns with an active ‘good-to-action’ transformation (Cai and Padoa-Schioppa, 2014).

Consequently, the preSMA’s role extends far beyond a passive motor relay, especially relevant to stimulus-to-action choices, it acts as an active across-domain transformer. In computational terms, while the vmPFC resolves the value of a stimulus, it lacks the mapping of this stimulus to the sensorimotor contingencies required to interact with it. The robust, sequential encoding of stimulus identity followed by action mapping within the preSMA suggests this region houses the transformational logic required to convert a chosen stimulus-value into a concrete behavioral policy. By holding both the stimulus identity and the motor effector, the preSMA dynamically routes the output of the prefrontal valuation network into the specific motor effectors required for goal acquisition, effectively acting as the central hub where value-based intention is translated into physical consequence.

Ultimately, our results provide a framework for understanding how the human frontal cortex bridges the gap between abstract valuation and concrete choice execution. We demonstrate that translating value into choice is not achieved through a single, static pathway. Instead, the brain leverages highly dynamic representations, relying on a shared set of neurons for the encoding of pre-decision values, before rapidly shifting into specialized, modality-specific roles to value and execute the finalized choice.

## Materials and Methods

### Participants

Sixteen adult patients (see Table S1 for demographics) with drug-resistant epilepsy participated in this study. All patients were undergoing stereotactic depth electrode implantation for seizure localization and potential surgical treatment at Cedars-Sinai Medical Center. Electrode placement was determined solely on clinical grounds by the clinical care team. Each patient provided written informed consent (Feinsinger et al., 2022) to participate in research protocols approved by the Institutional Review Boards of both the California Institute of Technology (IR23-1313) and Cedars-Sinai Medical Center (STUDY00000572).

### Task Design

#### General Experimental Structure

Participants (see Table S1 for demographics) performed two variants of a probabilistic reinforcement learning task: an Action Task and a Stimulus Task. Each trial followed a consistent temporal structure: a fixation cross (500–1500 ms), a response window (max 3000 ms), a selection confirmation (750 ms), and a final outcome phase (1250 ms). Rewards were represented by a yellow coin (value = 1), while non-rewarded trials were indicated by a red coin (value = 0). The order of these tasks was counterbalanced across sessions (Stimulus Task first: N=13 sessions). Reward probabilities were (re-)set at the start of each block (36 trials) and were pseudo-randomly drawn from the following set [*P* = 0.8*,P* = 0.6*,P* = 0.4*,P* = 0.2] such that each choice option was matched with an unique reward probability.

#### Action-Based Learning Task

In the action task, reward probabilities were mapped directly to specific motor actions (left hand, right hand, left foot, or right foot press). In each block, a subset of three actions was available for selection. Participants had to learn which action yielded the highest probability of reward over 36 trials. Participants completed between 4-12 blocks of the action task (5.7 blocks on average across sessions).

#### Stimulus-Based Learning Task

The stimulus task presented a binary choice between two stimuli drawn from a set of three available within a block (out of the total set of four; tree, house, dog, and camera). To decouple reward learning from motor responses, stimulus locations (top or bottom and left or right) were pseudo-randomized. Top locations required a hand press, while bottom locations required a foot press. Reward probability was dependent solely on the stimulus identity. Participants completed between 3-9 blocks of the stimulus task (5.25 blocks on average).

### Behavioral Data Analysis

#### Learning Curves and Choice Probability

To visualize behavioral adaptation over the course of the block, we generated per-subject learning curves by plotting the probability of selecting the highest-value option across trial indices. A linear regression was fitted to each individual subject’s choice data across the 36 trials to capture individual learning trajectories.

To determine if participants successfully learned the task parameters, we isolated behavioral performance during the final 10 trials (trials 27–36) of each block. Overall group performance was tested using a one-sided Wilcoxon signed-rank test, comparing the median accuracies of all subjects in the final 10 trials against chance.

#### Reward History and Choice Continuity (GLMs)

To quantify how recent reward history influenced choice perseveration, we implemented a combined-history logistic regression model. The model predicted the binary outcome of choice continuity (staying with the previous choice vs. switching) as a function of the rewards received on the preceding three trials (*t* – 1, *t –* 2, and *t –* 3).

The logistic regression was specified as:

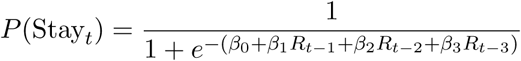

where *R_t–k_* represents the binary reward outcome at lag *k*. From the fitted model, we extracted the logit coefficients (*β*) to represent the log-odds of staying, as well as the marginal probability increase. The probability increase was calculated by taking the difference between the baseline probability of staying (derived from the intercept *β*_0_) and the probability of staying given a reward at a specific lag. Group-level significance for the effect of each lag was assessed by testing the subject-level coefficients against zero using a one-sample *t*-test.

### Computational Modeling

#### Reinforcement learning - 2 parameter

We modeled participant choice behavior using a standard reinforcement learning framework that relied solely on expected values (*Q*).

#### Choice Rule

The probability *P* of choosing option *i* on trial *t* was determined by a softmax choice rule:

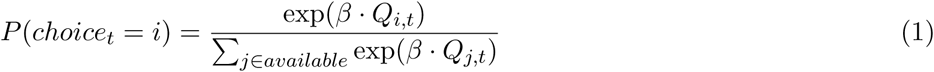

where *β* is the inverse temperature parameter governing choice stochasticity.

#### Update Rules

Q-values were updated based on the reward prediction error:

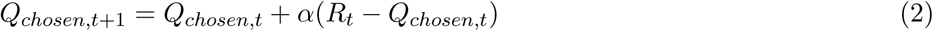

where *R_t_* is the reward and *α* is the learning rate.

#### Model Fitting

Parameters (*α, β*) were estimated by minimizing the regularized negative log-likelihood:

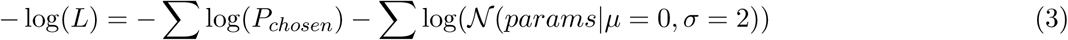

We utilized a normal distribution prior (*µ* = 0*, σ* = 2) to regularize the parameters during the optimization process.

### Choice kernel only model

We modeled participant choice behavior using a framework that relied entirely on choice history (*CK*), completely omitting reward-based value learning.

#### Choice Rule

The probability *P* of choosing option *i* on trial *t* was determined by a choice-history softmax rule:

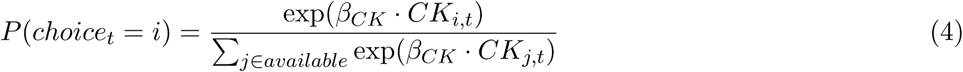

where *β_CK_* represents the strength of choice perseveration (or alternation, if negative).

#### Update Rules

The choice kernel was updated exclusively to track past choices:

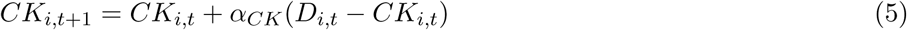

where *D_i,t_* is a dummy variable (1 if chosen, 0 otherwise) and *α_CK_* is the kernel learning rate (decay).

#### Model Fitting

Parameters (*α_CK_, β_CK_*) were estimated using the regularized negative log-likelihood optimization described in Equation 3.

### Reinforcement learning - 4 parameter

We modeled participant choice behavior using a hybrid reinforcement learning framework that integrated both expected values (*Q*) and choice history (*CK*).

#### Choice Rule

The probability *P* of choosing option *i* on trial *t* was determined by a combined softmax choice rule:

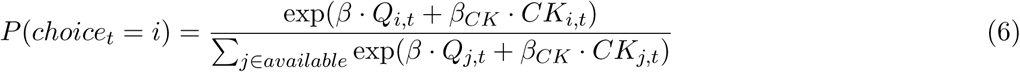

where *β* is the inverse temperature parameter and *β_CK_* represents the strength of choice perseveration.

#### Update Rules

Both systems operated in parallel: Q-values were updated based on reward prediction errors (Equation 2), and the choice kernel was simultaneously updated to track choice history (Equation 5).

#### Model Fitting

The full parameter set (*α, β, α_CK_, β_CK_*) was estimated utilizing the identical regularized optimization procedure established in Equation 3.

### Posterior Predictive Checks

#### Simulation Procedure

To evaluate how well our fitted models captured the empirical behavioral patterns, we conducted posterior predictive checks. For each participant, we generated synthetic data using their best-fitting parameters for three models: a standard Reinforcement Learning model (RL2; *α, β*), a Choice Kernel model (CK; *α_CK_, β_CK_*), and a hybrid model combining both (RL4; *α, β, α_CK_, β_CK_*).

To account for the stochastic nature of the models’ decision rules, we simulated 100 independent experimental runs per participant, per model, for both the Action and Stimulus tasks. Each simulation strictly mirrored the empirical task structure experienced by that specific participant, including the exact number of blocks, trials, and the specific trial-by-trial bandit configurations. To ensure reproducibility, simulation stochasticity was controlled using unique random seeds derived from participant identifiers.

#### Behavioral Metrics

We assessed model performance by comparing the synthetic data to the empirical data across key behavioral signatures of reinforcement learning:

- **Win-Stay / Lose-Shift Probabilities:** We calculated the raw probability of a participant repeating their previous choice, conditioned separately on whether the previous trial resulted in a reward or no reward. Trials where the previously chosen option was no longer available were excluded from this analysis.
- **Lagged Reward Effects:** To quantify the extended impact of past rewards on current choices, we fitted trial-by-trial logistic regression models predicting the probability of staying with the same choice. The model included the outcomes of the three most recent trials (lags *t –* 1, *t –* 2, and *t –* 3) as predictors.

#### Simulation Aggregation

For the simulated datasets, the behavioral metrics (stay probabilities and logistic regression estimates) were calculated independently for each of the 100 simulated runs. These values were then averaged within each participant to produce a single, stable model prediction per metric. These averaged predictions were directly compared against the metrics derived from the participants’ actual empirical data to evaluate model fit.

### Neural data

#### Electrophysiology and Recording

We used Behnke-Fried hybrid depth electrodes (AdTech Medical), positioned exclusively according to clinical criteria (Carlson et al., 2018)(see Supplementary Figure S1 for electrode locations). Broadband extracellular recordings were performed with a sampling rate of 32 kHz and a bandpass of 0.1–9,000 Hz (ATLAS System, Neuralynx). The dataset of 902 single units reported in this study was obtained bilaterally from the ventromedial prefrontal cortex (vmPFC), anterior cingulate cortex (ACC), pre-supplementary motor area (preSMA), and amygdala, with typically one macroelectrode targeted to each region per hemisphere. Each macroelectrode contained a bundle of eight 40-*µ*m microelectrodes extending past the macroelectrode tip. Recordings were bipolar, utilizing one microelectrode in each bundle as a local reference to isolate localized single-unit activity.

#### Spike Sorting and Unit Isolation

Spike sorting and unit isolation were performed offline using the open-source OSort algorithm (Rutishauser et al., 2006). Spike detection was performed using an amplitude threshold set to five standard deviations above the median noise level of the filtered signal.

Following detection, the OSort algorithm automatically clustered the extracted waveforms. This automated step grouped spikes into distinct putative units based on a distance metric utilizing their waveform features. After automatic clustering, all putative units underwent rigorous manual curation to ensure high data quality. We evaluated each cluster based on waveform shape consistency, temporal stability across the recording session, and the distribution of inter-spike intervals (ISIs).

To ensure the purity of our single-unit isolations, we strictly enforced a physiological refractory period. Any cluster exhibiting more than a 3% violation rate within a 3 ms ISI period was classified as multi-unit activity or noise and excluded from the single-unit analyses. Furthermore, automated clusters representing mechanical or electrical artifacts were manually identified and discarded. This combination of automated sorting and rigorous manual refinement yielded the final dataset of 902 high-quality single units utilized for all subsequent analyses.

#### Single-unit encoding analysis

To characterize how individual neurons represent value and choice identity, we modeled single-unit spike counts using Poisson generalized linear models (GLMs). For our primary epoch-based analyses, spike counts were extracted during fixed task windows, such as a 1000 ms window immediately following stimulus onset (for option value results) or a 1000 ms window preceding the execution of the motor response (for chosen value and chosen identity results).

GLM’s were constructed to included the variable of interest (e.g., chosen value, chosen identity) alongside relevant nuisance regressors (such as the availability of specific options on a given trial). Models were fit using Iteratively Reweighted Least Squares (IRLS); if convergence failed or algorithmic instability was detected, the optimizer automatically fell back to the L-BFGS algorithm.

The explanatory power of each variable was evaluated using a likelihood ratio test (LRT), comparing the log-likelihood of the full model to a reduced model omitting the specific term of interest (e.g. option values across the 4 stimuli). To robustly establish statistical significance and account for non-normal spike distributions, we computed empirical *p*-values via permutation testing. For each neuron, the variable of interest was randomly shuffled across trials 500 times. The empirical *p*-value was calculated as the proportion of permuted likelihood ratio statistics that met or exceeded the true, unpermuted statistic.

##### Proportion of significant units

To quantify whether a given brain region represented a variable of interest at the population level, we calculated the proportion of individually significant units within that region. A single unit was classified as significantly encoding the variable (e.g., chosen *Q*-value) if the *p*-value for the corresponding regressor in the first-level generalized linear model (GLM) fell below our predefined alpha level (*α* = 0.05). The total number of units included in the GLM for a specific region and task served as the denominator. To statistically determine if the observed proportion of significant units in a given region was greater than what would be expected by random chance, we performed a one-sided binomial test. The expected chance probability was mathematically tied to the single-unit significance threshold (*p* = 0.05). The resulting binomial *p*-values were evaluated at a significance threshold of *p <* 0.01.

##### Double-dissociation analysis

To formally test for functional specialization between regions across task domains (i.e., a double dissociation), we evaluated whether the likelihood of a neuron encoding chosen value was dependent on the interaction between brain region and task domain. We extracted the single-unit significance status (1 = significant at *α* = 0.05, 0 = not significant) for all units recorded in the predefined regions of interest (vmPFC and ACC) during both the stimulus and action tasks. Units included in the GLM for only one task were retained in the denominator for that specific task to ensure accurate, independent population proportions. We then fit a logistic regression model to predict the binary significance status of each unit. The model included categorical predictors for Brain Region (vmPFC vs. ACC), Task Domain (Stimulus vs. Action), and their interaction term (Region × Task). The formal test for the double dissociation was evaluated based on the statistical significance of the Region × Task interaction coefficient.

##### Sliding-window encoding analysis

To capture the fine-grained temporal dynamics of value and identity representations, we extended our GLM framework using a sliding-window approach. For each trial, spike counts were extracted within 500 ms rolling windows, advancing in 50 ms increments. These sliding windows were aligned to either the onset of the stimulus (spanning −500 to 1000 ms) or the execution of the motor response (spanning −1000 to 500 ms). The exact same joint Poisson GLM, optimization, and permutation testing procedures described above were applied independently to each time window. Units that survived the permutation at *p <* 0.05 are included in the proportion.

##### Latency analysis

To determine the precise timing of neural responses relative to task events, we computed response latencies for individual single units. This analysis was restricted to the subpopulation of units that demonstrated significant task-related encoding (*p <* 0.05, determined via the permutation-tested generalized linear models described above) to ensure latencies were only calculated for functionally responsive neurons.

Latency was defined as the first point in time at which a neuron’s spiking rate significantly deviated from its expected baseline firing rate. We estimated this baseline rate using a 500 ms window immediately preceding stimulus onset. To maximize robustness, particularly for neurons with sparse baseline firing, we calculated a session-wide baseline firing rate for each neuron by aggregating all pre-stimulus baseline spikes across the entire recording session. For each trial, we iteratively scanned the spike train relative to the event of interest (e.g., motor response).

Moving temporally away from the event, we tallied the cumulative number of spikes and calculated the exact time elapsed for each incoming spike. At each spike time, we computed the expected number of spikes based on the session-wide baseline Poisson rate. We then utilized the Poisson cumulative distribution function (CDF) and survival function (SF) to test whether the observed cumulative spike count represented a statistically significant deviation (*p <* 0.05) from the baseline expectation.

A significant increase in firing (a burst) was identified if the probability of observing the tallied number of spikes (or more) was less than 0.05. Conversely, a significant decrease in firing (a suppression) was identified if the probability of observing the tallied number of spikes (or fewer) fell below 0.05. The exact time elapsed to this first significant deviation was recorded as the trial’s response latency.

To compare the temporal dynamics of these responses across brain regions and task domains, we aggregated the significant trial-level latencies for each region. Differences in the median latency distributions between specific regions, as well as between the stimulus-based and action-based tasks within a given region, were evaluated for statistical significance using two-sided non-parametric Wilcoxon rank-sum tests.

#### Decoding generalization analysis

To determine whether representations of choice identity generalize across decision domains, we performed a cross-task population decoding analysis. For each brain region, we constructed pseudo-populations by pooling independently recorded single units across all participants. Spike counts for each unit were extracted during the window of interest and Z-scored. To prevent class imbalances from biasing the decoder, we randomly subsampled trials to ensure an equal number of observations per condition. Pseudo-population feature matrices were standardized prior to model fitting.

Since chosen option identity is a categorical variables, we utilized a Linear Support Vector Classifier (SVC) with L2 regularization. To explicitly test for representational generalization, each model was trained exclusively on data from one task domain (e.g., the action-based task) and tested on unseen data from the alternative domain (e.g., the stimulus-based task). Model performance was quantified using classification accuracy.

Statistical significance was established using a rigorous permutation and bootstrapping approach. First, to account for variance introduced by the random trial subsampling, the entire pseudo-population generation and cross-task testing pipeline was repeated across 50 independent random seeds. Within each seed, we generated a local null distribution by shuffling the test labels 100 times. To compute the final overarching significance, we employed a “null of means” bootstrapping procedure: we drew one permuted score from each of the 50 seeds to calculate a null mean, repeating this process 5,000 times. The empirical *p*-value was then calculated as the proportion of these bootstrapped null means that performed better than or equal to the true observed mean across the 50 seeds (i.e., higher accuracy for categorical variables).

## Acknowledgments

We would like to thank all members of the O’Doherty and Rutishauser laboratories for their feedback. We thank the participants and their families for their participation, and nurses and medical staff for their work. We would like to acknowledge funding from the following grants; NIH MH133729 and NSF 2318899 to J.P.D. and U.R., as well as NIH U01NS117839 to U.R.

