## Supplemental information for "Choice-driven remapping of action- and stimulus-anchored value in human single neurons"

### Supplementary Information

| Patient ID | Sex | Age | Ethnicity | Race |
|---|---|---|---|---|
| P97CS | M | 32 | Non-Hispanic | Black |
| P94CS | F | 26 | Hispanic | White |
| P92CS | F | 30 | Non-Hispanic | White |
| P90CS | M | 32 | Non-Hispanic | Black |
| P87CS | F | 26 | Hispanic | White |
| P86CS | F | 39 | Hispanic | White |
| P84CS | M | 63 | Non-Hispanic | White |
| P81CS | F | 28 | Hispanic | Other |
| P98CS | F | 30 | Hispanic | White |
| P99CS | F | 39 | Hispanic | White |
| P100CS | M | 25 | Hispanic | White |
| P103CS | M | 53 | Non-Hispanic | White |
| P102CS | F | 55 | Non-Hispanic | White |
| P106CS | M | 37 | Non-Hispanic | White |
| P108CS | F | 44 | Non-Hispanic | White |
| P110CS | M | 38 | Hispanic | White |

**Table S1:** Patient demographics.

| | Action Task | Stimulus Task |
|---|---|---|
| Choice Kernel | 7426.4 | 4541.0 |
| RL 2 parameter | 7464.5 | 4429.5 |
| RL 4 parameter | 6757.6 | 4444.3 |

**Table S2:** Total BIC scores across subjects.

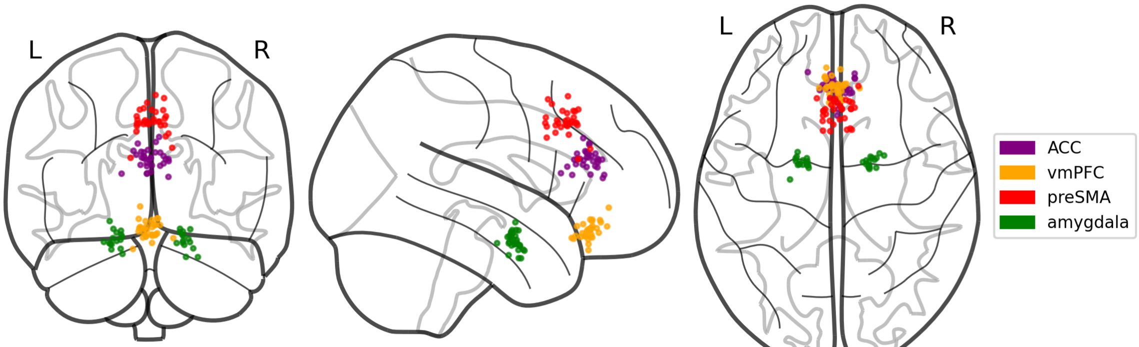


**Figure S1:** Locations of ACC, vmPFC, preSMA and amygdala electrodes across patients.

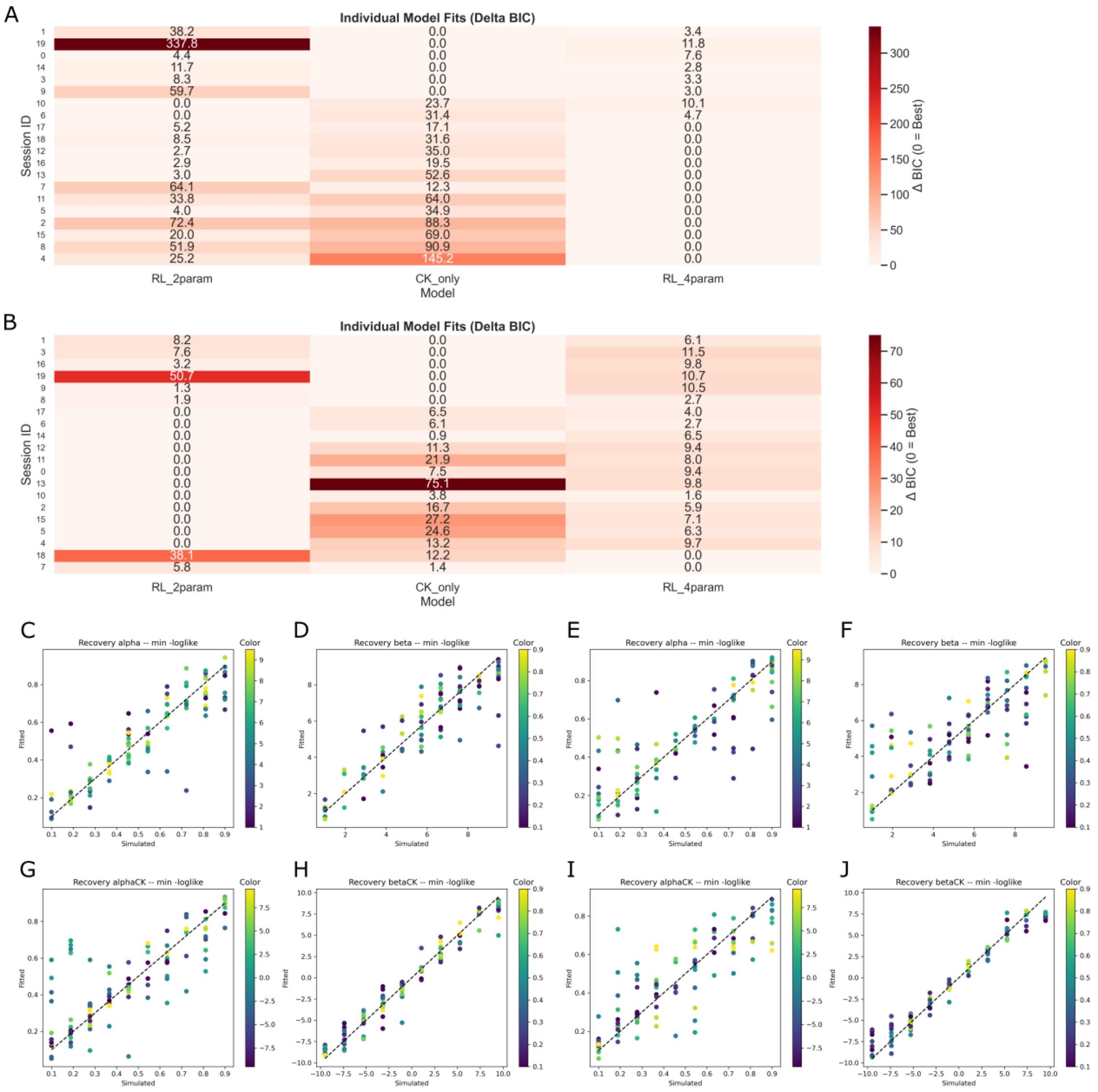


**Figure S2:** **A)** Individual session's model performance for the Action Task. Distance in BIC between the best (0 distance) and other models. Larger BIC distance indicates worse model performance. **B)** Similarly, Stimulus Task individual session's model performance. **C), D), G), H)** 4 param RL model parameter recovery in the stimulus task. **E), F), I), J)** 4 param RL model parameter recovery in action task.

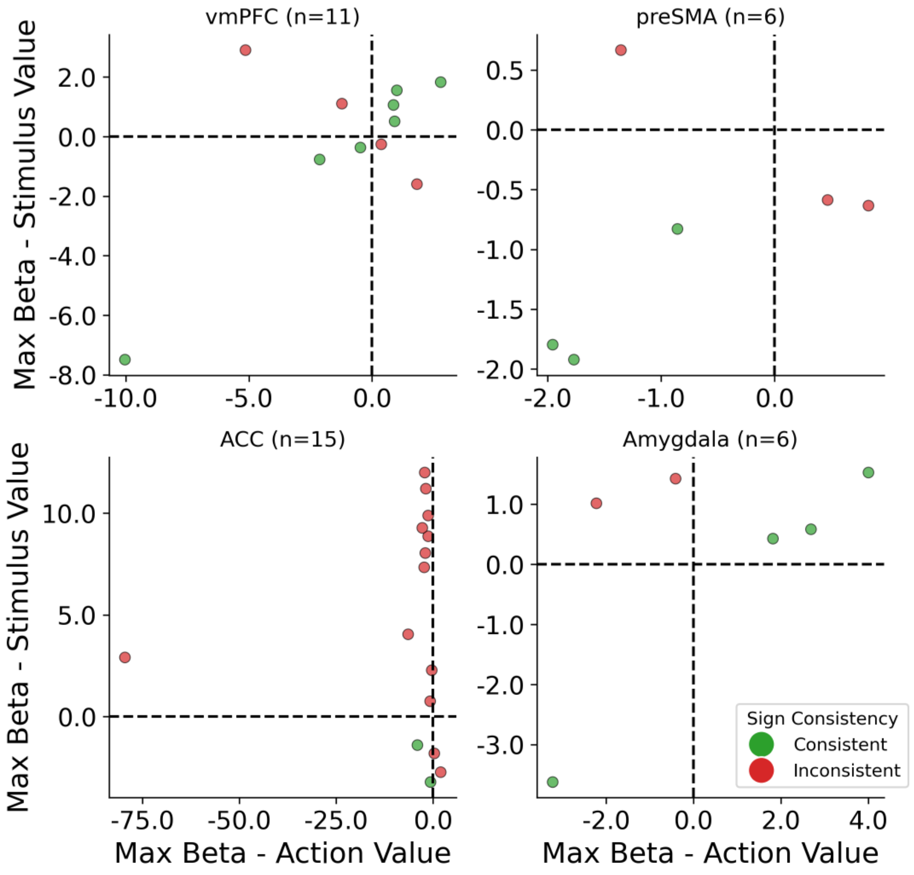


**Figure S3:** Consistency in coding direction of the largest beta. Neurons are selected to only include those that significantly ($p > 0.05$) code for both stimulus value in the stimulus task and action value in the action task. Model fitted beta coefficients with the largest effect (i.e., the strongest stimulus- or action-value code)

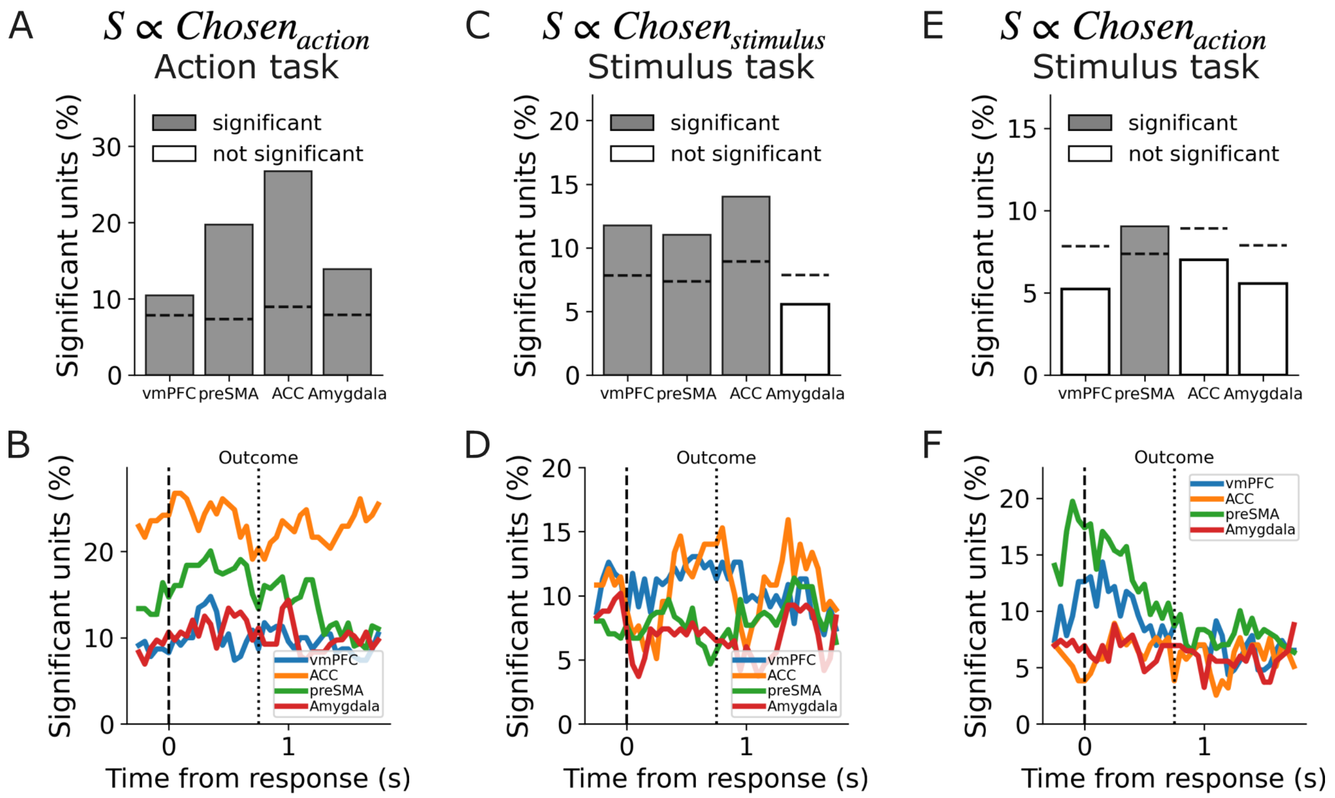


**Figure S4:** **A)** Proportion of significant neurons encoding the chosen action during the action task across four regions of interest: ventromedial prefrontal cortex (vmPFC), pre-supplementary motor area (preSMA), anterior cingulate cortex (ACC), and Amygdala. Color conventions match main manuscript **B)** Time-resolved proportion of significant units encoding the chosen action relative to response execution (dashed vertical line at 0 ms) in the action task. Solid lines denote the proportion for each brain region computed using a sliding window approach (see Methods). **C)** Proportion of significant neurons encoding the chosen stimulus during the stimulus task. Conventions follow panel A. **D)** Time-resolved proportion of significant units encoding the chosen stimulus relative to the response during the stimulus task. Conventions follow panel B. **E)** Proportion of significant neurons encoding the chosen action during the stimulus task. Conventions follow panel A. **F)** Time-resolved proportion of significant units encoding the chosen action relative to the response during the stimulus task. Conventions follow panel B
